# Defining critical roles for ZBP1 in PANoptosis utilizing a novel genetic tool for disease modeling and therapeutic development

**DOI:** 10.64898/2026.05.19.726380

**Authors:** Alexander P. Young, Twinu Wilson Chirayath, Yaqiu Wang, Sangappa B. Chadchan, Thirumala-Devi Kanneganti

**Affiliations:** Department of Immunology, St. Jude Children’s Research Hospital, Memphis, TN, 38105, USA

## Abstract

Innate immunity and innate immune cell death provide a critical first line of defense against disease. However, excess cell death leads to pathological inflammation. ZBP1 is an innate immune sensor that is central to this balance between defense and inflammation as a driver of inflammatory lytic cell death, PANoptosis. Activation of ZBP1-dependent PANoptosis downstream of diverse triggers has roles in both host defense and disease pathology, making ZBP1 an attractive therapeutic target. Therefore, understanding the distinct roles of ZBP1 in different cell types, organ systems, and tissues is critical to identify therapeutic strategies. Although ZBP1 regulates PANoptosis in multiple cell types, there are limited tools to interrogate its function in a cell type-specific manner. Here, we report the generation of a *Zbp1*-floxed mouse line (*Zbp1*^fl/fl^) for investigation of ZBP1 in distinct cell populations. We crossed *Zbp1*^fl/fl^ mice to *LysM*^cre^ mice to selectively deplete *Zbp1* from the myeloid compartment, which did not alter immune homeostasis. Bone marrow-derived macrophages (BMDMs) from *Zbp1*^fl/fl^ mice had normal ZBP1 expression and PANoptosis activation, while those from *Zbp1*^fl/fl^*LysM*^cre^ mice exhibited markedly reduced ZBP1 expression and were biochemically and functionally protected from ZBP1-driven PANoptosis; these effects were validated using known triggers of the ZBP1-PANoptosome—IAV, nuclear export inhibition plus IFN, and ethanol. These findings demonstrate this new *Zbp1*^fl/fl^ mouse as a versatile tool that can be utilized with a variety of Cre-drivers to study ZBP1 in a wide array of distinct cell types. Given the critical role of ZBP1 in disease, this tool will inform the development of therapeutic strategies.

## Introduction

The innate immune system provides the first line of defense against pathogen infection, oncogenesis, and altered homeostasis. Cell death is a key effector mechanism during innate immune responses to remove compromised cells and propagate inflammatory signaling. PANoptosis is an innate immune, lytic cell death pathway that is initiated by innate immune sensors and driven by caspases and RIPKs through PANoptosome complexes^1–7^. The innate immune sensor Z-DNA binding protein 1 (ZBP1) was the first PANoptosome sensor characterized, where it was described as a sensor of influenza A virus (IAV)^8^. ZBP1 was later found to drive PANoptosis in response to host nucleic acids in the absence of ADAR1. This pathway can be therapeutically stimulated by IFN in conjunction with nuclear export inhibitors (NEI; e.g., KPT-330) to regress tumors in murine models^9^. Subsequent studies have identified numerous other infectious and sterile triggers that drive ZBP1-dependent PANoptosis with implications across disease^10–13^.

ZBP1-dependent PANoptosis has been best characterized in primary bone marrow-derived macrophages (BMDMs), but current evidence suggests that ZBP1 also drives PANoptosis across other cell types. For example, mouse hepatocytes undergo ZBP1-dependent PANoptosis in response to ethanol, similar to BMDMs^12^. ZBP1 also drives PANoptosis in a variety of human and mouse endothelial cells^14,15^, fibroblasts^16–18^, and central nervous system cell types^19,20^. Furthermore, several types of cancer cells have demonstrated susceptibility to ZBP1-mediated PANoptosis^13,21^. Collectively, the growing ZBP1 literature has defined its roles, as well as the roles of ZBP1-dependent PANoptosis, in infectious disease, inflammatory disorders, neurodegenerative disease, and cancer^9–11,22,23^.

Although ZBP1 regulates inflammatory cell death in a variety of cell types in both mice and humans, its cell type-specific functions are not well characterized *in vivo*, and there is a lack of biological tools to investigate ZBP1 in specific cell types of interest. Therefore, in this study, we developed a *Zbp1*-floxed mouse (*Zbp1*^fl/fl^) for conditional deletion of *Zbp1* to interrogate the function of ZBP1 in Cre-Lox compatible cell types. Using *LysM*^cre^ as a proof-of-concept Cre driver, we selectively depleted ZBP1 from the myeloid compartment. Myeloid ZBP1 depletion did not impact numbers of myeloid or other immune cells, indicating that the insertion of loxP sites or recombination with *LysM*^cre^ did not impact immune homeostasis. Furthermore, BMDMs from *Zbp1*^fl/*fl*^*LysM*^*cre*^ mice exhibited markedly reduced ZBP1 expression and were biochemically and functionally protected from ZBP1-dependent PANoptosis in response to IAV infection, KPT-330 plus IFN-γ, or ethanol. Therefore, this validated *Zbp1*^fl/fl^ mouse line can be used to systematically investigate the functions of ZBP1 in a wide range of cell types using different Cre drivers. This genetic tool will aid the study of ZBP1 in a cell-specific manner in the numerous disease conditions where ZBP1 plays a role and inform the development of therapeutic strategies.

## Results

### Design and generation of *Zbp1*^fl/fl^ mice

ZBP1 is a critical innate immune sensor that uses its Zα domains to promote inflammatory and cell death signaling^1,8,24,25^. To generate *Zbp1*^fl/fl^ mice, loxP sites were inserted flanking exons 2 and 3 to delete the Zα domains and create a functional knockout in mice containing a transgenic Cre driver (Figure 1A). Genotyping of the 5’ junction (Figure 1B) and the 3’ junction (Figure 1C) confirmed the presence of the loxP sites in *Zbp1*^fl^ mice. We then used these *Zbp1*^fl/fl^ mice to generate myeloid-specific *Zbp1* conditional knockout mice by crossing them to *LysM*^cre^ mice. This yielded *Zbp1*^fl/fl^*LysM*^cre^ mice and *Zbp1*^fl/fl^ littermate controls which are expected to phenocopy wild-type mice. As mice that are homozygous for the *LysM*^cre^ allele would effectively become *Lyz2* knockouts, only *Zbp1*^fl/fl^*LysM*^cre^ mice that were heterozygous for *LysM*^cre^ were used.

**Figure 1.**
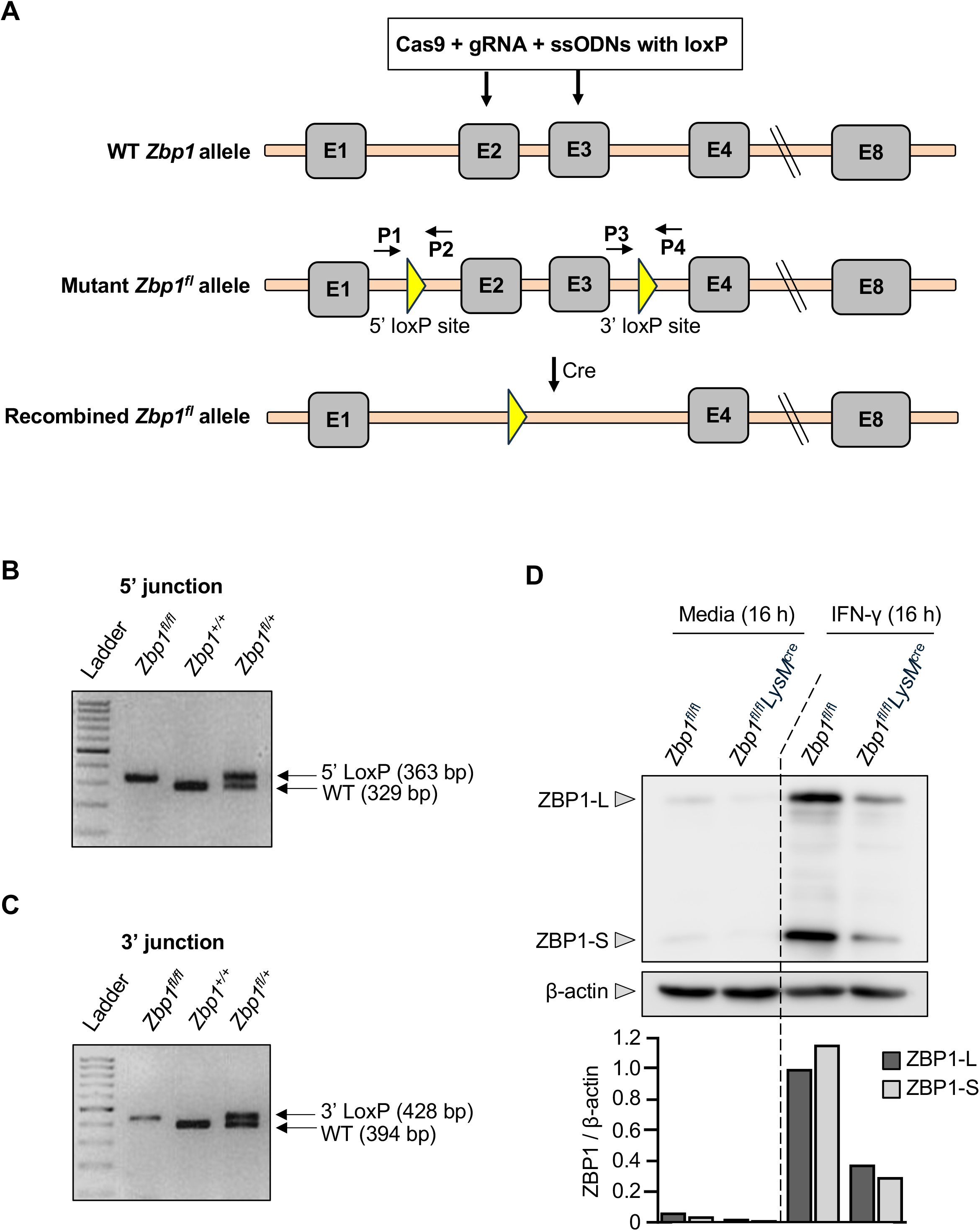
Design of *Zbp1*^fl/fl^*LysM*^cre^ mice. **A**) Schematic representation of the genetic strategy used to generate a mouse line carrying a floxed allele of *Zbp1*. Cas9, guide RNAs (gRNA), and single-stranded oligodeoxynucleotides (ssODNs) were utilized to insert loxP sites flanking exon 2 and exon 3 of the *Zbp1* allele. **B, C**) Representative PCR-based genotyping of *Zbp1*^fl/fl^, *Zbp1*^+/+^, and *Zbp1*^fl/+^, mice confirming insertion of the loxP sites in the 5’ junction (**B**) and 3’ junction (**C**) of the *Zbp1* locus. **D**) BMDMs from *Zbp1*^fl/fl^ or *Zbp1*^fl/fl^*LysM*^cre^ mice were treated with IFN-γ or media control. Representative immunoblot (top) and quantification of ZBP1 relative to β-actin by densitometric analyses at 16 h, demonstrating the abundance of ZBP1 long (ZBP1-L) or short (ZBP1-S) isoforms. The abundance of ZBP1 was normalized to the ZBP1-L isoform in *Zbp1*^fl/fl^ cells treated with IFN-γ.

Both isoforms of ZBP1, the long isoform (ZBP1-L) and short isoform (ZBP1-S), are relevant to the overall role of ZBP1 in inflammation and cell death^26,27^. Stimulation with IFN-γ induced the expression of both ZBP1-L and ZBP1-S in *Zbp1*^fl/fl^ BMDMs, as expected (Figure 1D). Meanwhile, BMDMs from *Zbp1*^fl/fl^*LysM*^cre^ mice displayed a > 60% depletion of both ZBP1-L and ZBP1-S under IFN-γ stimulation (Figure 1D). Together, these data confirmed the successful generation of *Zbp1*^fl/fl^*LysM*^cre^ mice, where ZBP1 expression is reduced in BMDMs.

### *Zbp1*^fl/fl^*LysM*^cre^ mice have similar abundances of immune cells compared to controls

Using these newly generated *Zbp1*^fl/fl^*LysM*^cre^ mice, we assessed the abundance of different immune cells to confirm that the addition of loxP sites to the *Zbp1* alleles and/or recombination with *LysM*^cre^ would not alter different immune cell compartments. To this end, we performed flow cytometry on isolated spleen and peritoneal cells from *Zbp1*^fl/fl^*LysM*^cre^ and control mice. Among splenocytes, we saw no difference between *Zbp1*^fl/fl^*LysM*^cre^ mice and the controls with respect to the abundance of CD4^+^ T cells, CD8^+^ T cells, natural killer (NK) cells, eosinophils, neutrophils, B cells, or macrophages (Figure 2A). Among peritoneal lavage cells, we saw no difference between *Zbp1*^fl/fl^*LysM*^cre^ mice and the controls in any of these cellular populations as well as no difference in the abundance of mast cells or monocytes (Figure 2B). These data indicate that depletion of ZBP1 specifically from the myeloid compartment did not affect the abundance of myeloid cells (e.g., monocytes, macrophages, or granulocytes) or other immune cell types (e.g., lymphoid cells). These results suggest that immune homeostasis is not overtly impacted in *Zbp1*^fl/fl^*LysM*^cre^ mice.

**Figure 2.**
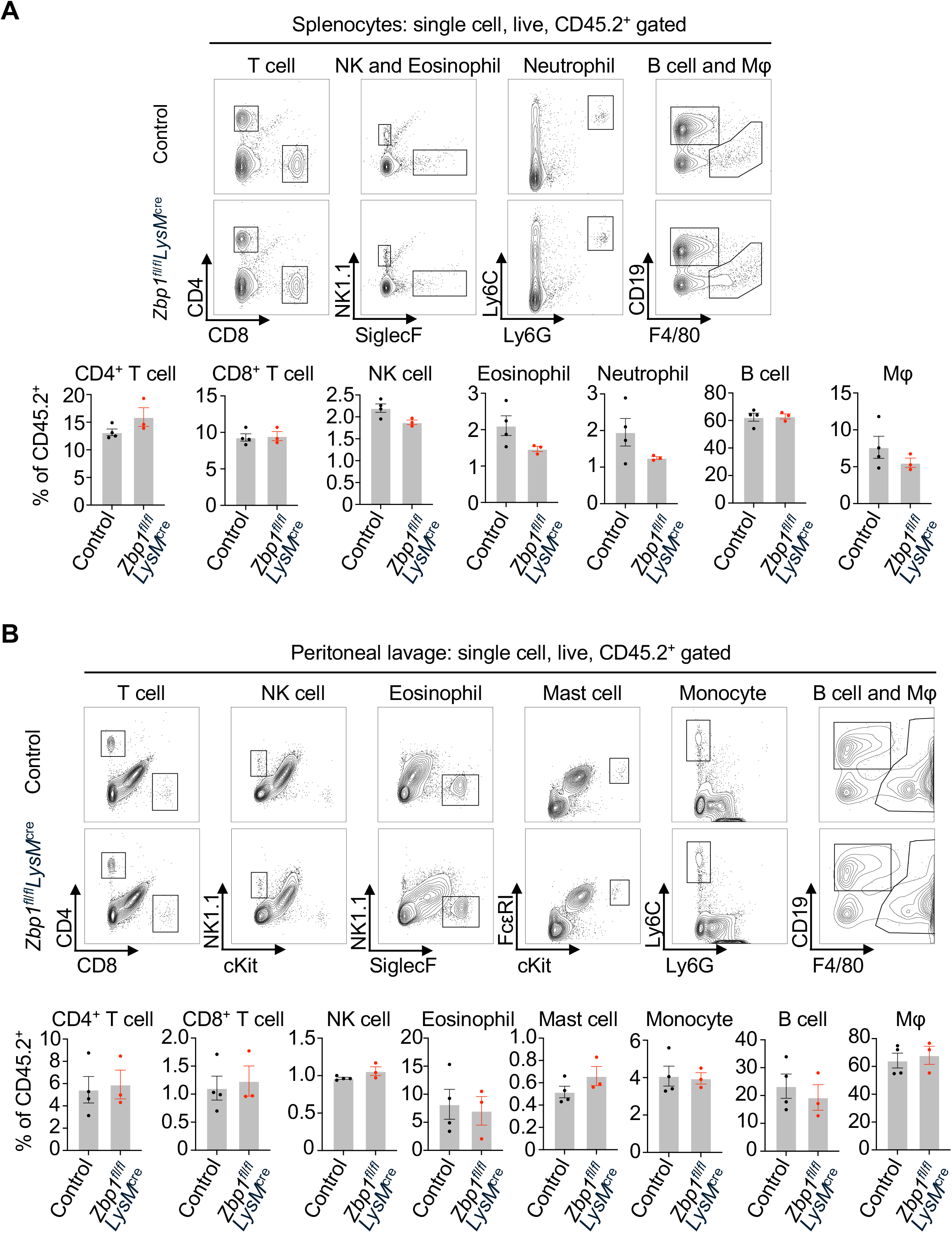
*Zbp1*^fl/fl^*LysM*^cre^ mice have similar abundances of myeloid and other immune cells compared to control mice. **A**) Total splenocytes were isolated from control (a pool of wild-type, *Zbp1*^fl/fl^, and *Zbp1*^+/+^*LysM*^cre^ mice) and *Zbp1*^fl/fl^*LysM*^cre^ mice. Flow cytometry was used to assess the relative abundance of CD4^+^ T cells, CD8^+^ T cells, natural killer (NK) cells (NK1.1^+^), eosinophils (SiglecF^+^), neutrophils (Ly6C^+^Ly6G^+^), B cells (CD19^+^), and macrophages (F4/80^+^) after gating for single, live (DAPI^-^), CD45^+^ cells. **B**) Total intraperitoneal cells were collected via peritoneal lavage from control (a pool of wild-type, *Zbp1*^fl/fl^, and *Zbp1*^+/+^*LysM*^cre^ mice) and *Zbp1*^fl/fl^*LysM*^cre^ mice. Flow cytometry was used to assess the relative abundance of CD4^+^ T cells, CD8^+^ T cells, NK cells (NK1.1^+^), eosinophils (SiglecF^+^), mast cells (FcεRI^+^cKit^+^), monocytes (Ly6C^+^Ly6G^-^), B cells (CD19^+^), and macrophages (F4/80^+^) after gating for single, live (DAPI^-^), CD45^+^ cells. Data are representative of 3–4 biological replicates from one independent experiment. Data are presented as mean ± SEM.

### *Zbp1*^fl/fl^*LysM*^cre^ BMDMs are protected from IAV-induced PANoptosis

We next aimed to assess whether the reduction in ZBP1 expression in the *Zbp1*^fl/fl^*LysM*^cre^ mouse line could prevent PANoptosis. To this end, we infected BMDMs from WT, *Zbp1*^fl/fl^*LysM*^cre^, and *Zbp1*^fl/f^ littermate control mice with IAV (MOI 25) to induce ZBP1-dependent PANoptosis. BMDMs from *Zbp1*^−/−^ mice were used as a negative control. IAV induced robust cell death among all wild-type and *Zbp1*^fl/fl^ BMDMs, reaching 100% by 16 h post-infection (Figure 3A–B). Cell death was reduced to 20% in *Zbp1*^fl/fl^*LysM*^cre^ BMDMs, and minimal death was observed in *Zbp1*^−/−^ BMDMs (Figure 3A–B).

**Figure 3.**
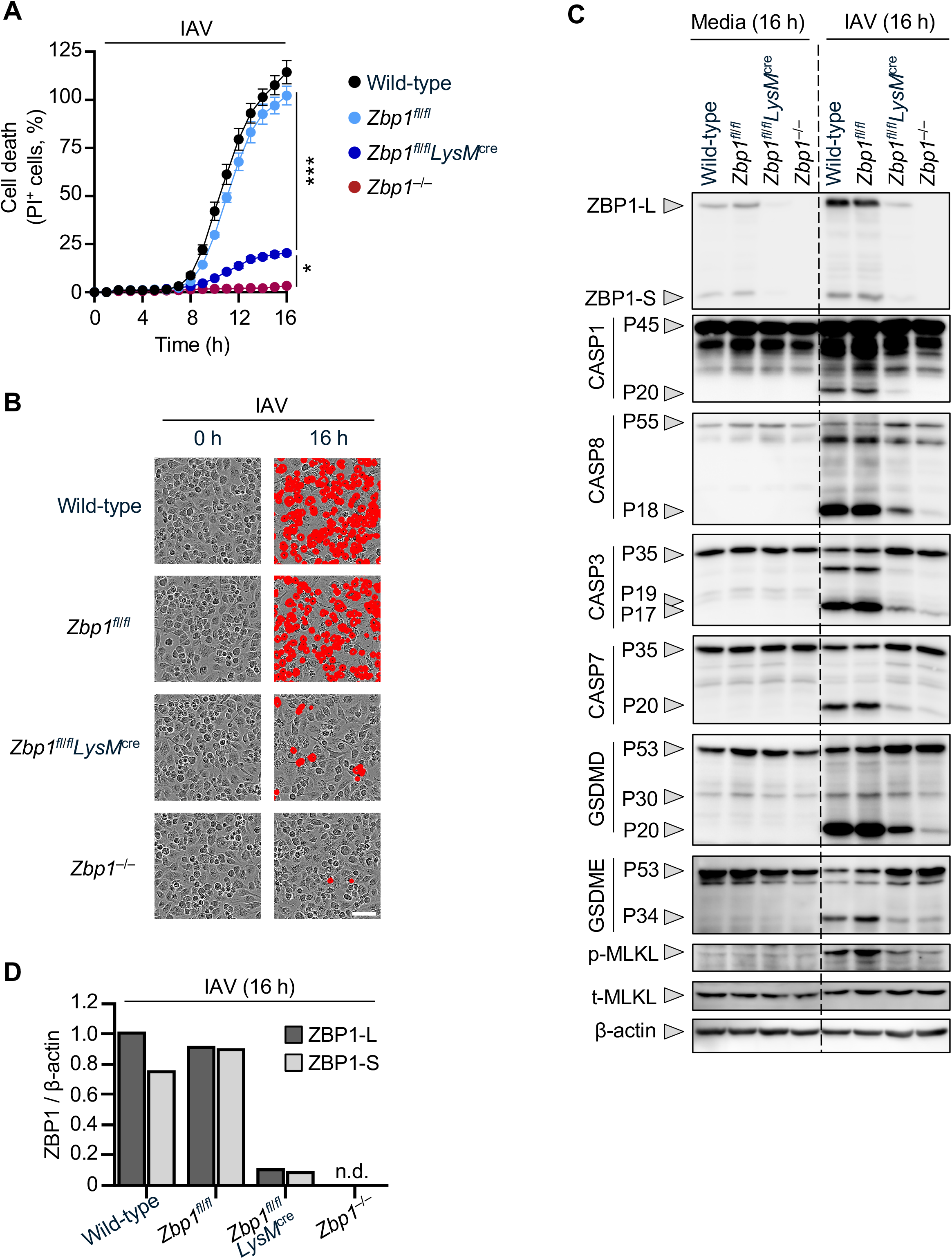
*Zbp1*^fl/fl^*LysM*^cre^ BMDMs undergo reduced PANoptosis in response to IAV infection. **A**) Cell death kinetics as assessed by PI uptake in wild-type, *Zbp1*^fl/fl^, *Zbp1*^fl/fl^*LysM*^cre^, and *Zbp1*^−/−^ BMDMs during influenza A virus (IAV) infection (MOI 25) over 16 h. **B**) Representative live-cell images captured at 0 h and 16 h during IAV infection. Cells that were determined to be PI^+^ are masked in red. **C**) Assessment of cell death proteins following treatment with media or infection with IAV for 16 h. Immunoblot was used to assess ZBP1 in the long (ZBP1-L) and short (ZBP1-S) isoform, pro-(P45) and active (P20) caspase-1 (CASP1), pro- (P55) and cleaved (P18) caspase-8 (CASP8), pro- (P35) and cleaved (P19/P17) caspase-3 (CASP3), pro- (P35) and cleaved (P20) caspase-7 (CASP7), pro- (P53), active (P30), and inactive (P20) gasdermin D (GSDMD), pro- (P53) and active (P34) gasdermin E (GSDME), phosphorylated MLKL (p-MLKL), and total MLKL (t-MLKL). β-actin was used as a loading control. **D**) ZBP1-L and ZBP1-S proteins were quantified by densitometry relative to β-actin, using the blots presented in panel C. The abundance of ZBP1 was normalized to the full-length (ZBP1-L) isoform in wild-type cells. Data are representative of at least three independent biological replicates (**A**–**D**). Data are presented as mean ± SEM and comprise four technical replicates from one experiment (**A**). Scale bar, 50 μm (**B**). Significance was determined by two-way ANOVA (**A**). * *p* < 0.05, *** *p* < 0.001.

To confirm that the resistance of *Zbp1*^fl/fl^*LysM*^cre^ BMDMs to cell death was consistent with reduced PANoptosis, we assessed the activation of key PANoptotic molecules, including caspases-1/8/3/7 and the pore-forming executioners, GSDMD, GSDME, and MLKL^28^. At 16 h post-infection, wild-type BMDMs exhibited upregulation of ZBP1, activation of all caspases, and activation of executioner molecules (i.e., cleavage of GSDMD and GSDME, and phosphorylation of MLKL; Figure 3C). The activation of these proteins in *Zbp1*^fl/fl^ BMDMs was comparable to that of wild-type cells (Figure 3C). In contrast, ZBP1 induction was markedly reduced in *Zbp1*^fl/fl^*LysM*^cre^ BMDMs during IAV infection, though some residual ZBP1 expression was detected (Figure 3C–D). Correspondingly, there was a clear reduction in the activation of each of the PANoptotic molecules in *Zbp1*^fl/fl^*LysM*^cre^ BMDMs, with some caspase and executioner activation still present. These biochemical results are consistent with the low amount of cell death observed in *Zbp1*^fl/fl^*LysM*^cre^ BMDMs (Figure 3A–B). Thus, the small degree of cell death and the activation of cell death proteins in *Zbp1*^fl/fl^*LysM*^cre^ BMDMs was commensurate with the residual expression of ZBP1. These data collectively indicate that the insertion of loxP sites does not impact ZBP1 cell death effector function in *Zbp1*^fl/fl^ BMDMs and that the reduction of ZBP1 in *Zbp1*^fl/fl^*LysM*^cre^ BMDMs (Figure 1C–D, Figure 3C–D) is sufficient to suppress PANoptosis in infected cells.

### *Zbp1*^fl/fl^*LysM*^cre^ BMDMs are protected from NEI plus IFN-γ-induced PANoptosis

We next sought to extend our findings to a model of ZBP1-driven PANoptosis independent of infection. We induced ZBP1-dependent PANoptosis by combined treatment with a NEI (KPT-330) plus IFN-γ^9^. Treatment with KPT-330 plus IFN-γ induced robust cell death in wild-type and *Zbp1*^fl/fl^ BMDMs (72% and 76%, respectively; Figure 4A–B), while the cell death was reduced to 50% in *Zbp1*^fl/fl^*LysM*^cre^ BMDMs and 31% in *Zbp1*^−/−^ cells by 24 h post-treatment (Figure 4A–B).

**Figure 4.**
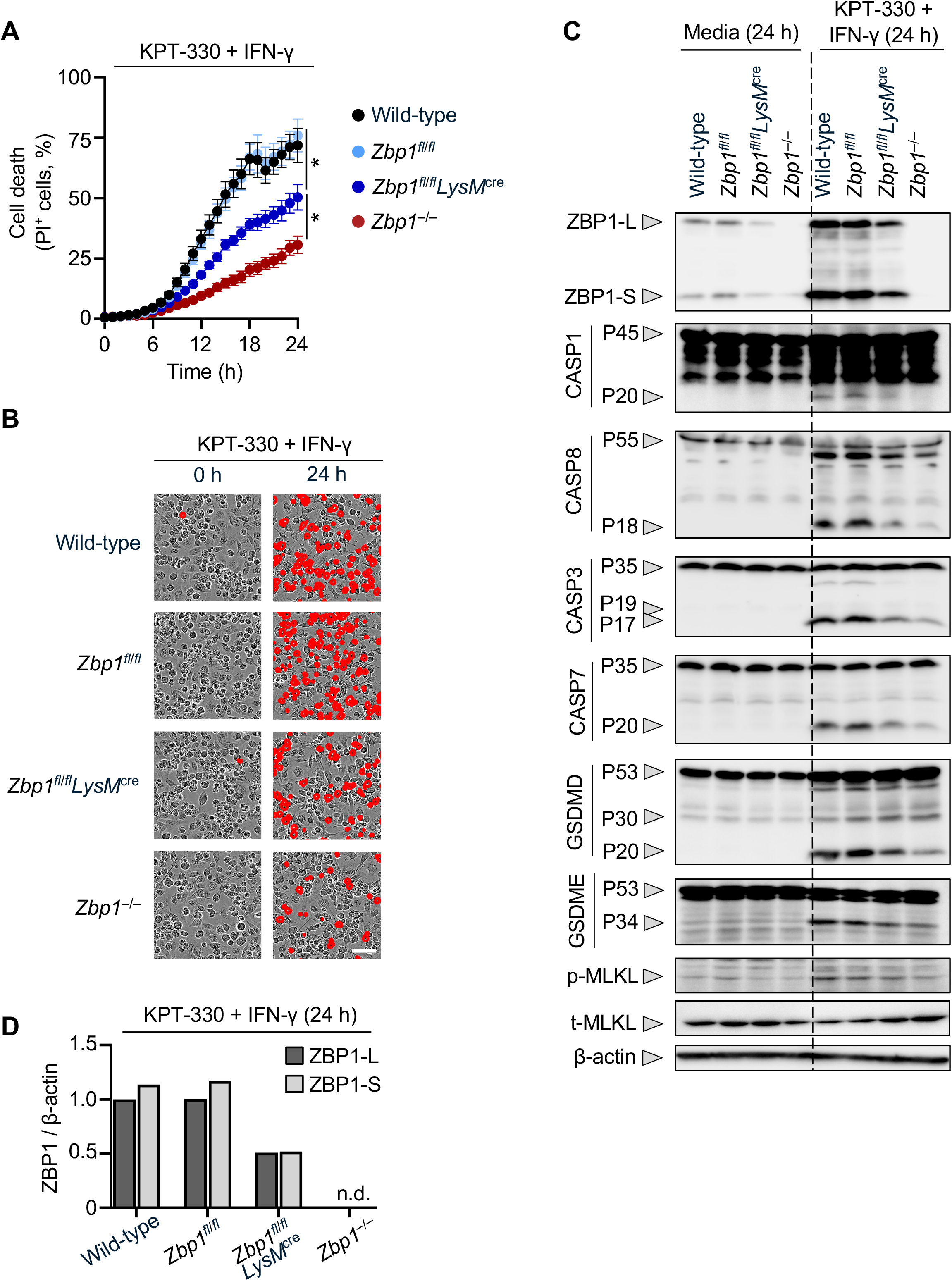
*Zbp1*^fl/fl^*LysM*^cre^ BMDMs undergo reduced PANoptosis in response to KPT-330 plus IFN-γ. **A**) Cell death kinetics as assessed by PI uptake in wild-type, *Zbp1*^fl/fl^, *Zbp1*^fl/fl^*LysM*^cre^, and *Zbp1*^−/−^ BMDMs during treatment with KPT-330 (5 μM) plus IFN-γ (50 ng/mL) over 24 h. **B**) Representative live-cell images captured at 0 h and 24 h following treatment with KPT-330 and IFN-γ. Cells that were determined to be PI^+^ are masked in red. **C**) Assessment of cell death proteins following treatment with media or KPT-330 plus IFN-γ for 24 h. Immunoblot was used to assess ZBP1 in the long (ZBP1-L) and short (ZBP1-S) isoform, pro- (P45) and active (P20) caspase-1 (CASP1), pro- (P55) and cleaved (P18) caspase-8 (CASP8), pro- (P35) and cleaved (P19/P17) caspase-3 (CASP3), pro- (P35) and cleaved (P20) caspase-7 (CASP7), pro- (P53), active (P30), and inactive (P20) gasdermin D (GSDMD), pro- (P53) and active (P34) gasdermin E (GSDME), phosphorylated MLKL (p-MLKL), and total MLKL (t-MLKL). β-actin was used as a loading control. **D**) ZBP1-L and ZBP1-S proteins were quantified by densitometry relative to β-actin, using the blots presented in panel C. The abundance of ZBP1 was normalized to the full-length (ZBP1-L) isoform in wild-type cells. Data are representative of at least three independent biological replicates (**A**–**D**). Data are presented as mean ± SEM and comprise four technical replicates from one experiment (**A**). Scale bar, 50 μm (**B**). Significance was determined by two-way ANOVA (**A**). * *p* < 0.05.

The activation status of PANoptotic proteins in response to KPT-330 plus IFN-γ exhibited a similar pattern as with IAV (Figure 4C). At 24 h post-treatment, wild-type and *Zbp1*^fl/fl^ BMDMs exhibited activation of caspases-1/8/3/7, cleavage of GSDMD and GSDME, and phosphorylation of MLKL (Figure 4C). Meanwhile, activation of these proteins was reduced in *Zbp1*^fl/fl^*LysM*^cre^ BMDMs, with a further reduction seen in *Zbp1*^−/−^ BMDMs (Figure 4C). ZBP1 protein levels were reduced but still detected in *Zbp1*^fl/fl^*LysM*^cre^ BMDMs (Figure 4C–D). Thus, the residual activation of PANoptosis signaling and low levels of cell death in *Zbp1*^fl/fl^*LysM*^cre^ BMDMs was correlated with the presence of residual ZBP1. These data together indicate that the reduction in ZBP1 levels in *Zbp1*^fl/fl^*LysM*^cre^ BMDMs is sufficient to significantly suppress ZBP1-dependent PANoptosis induced by KPT-330 plus IFN-γ.

### *Zbp1*^fl/fl^*LysM*^cre^ BMDMs are resistant to ethanol-induced PANoptosis

To further assess the utility of the *Zbp1*^fl/fl^*LysM*^cre^ system, we utilized ethanol, another trigger of ZBP1-dependent PANoptosis^12^. Ethanol (5% v/v) treatment induced robust cell death in both wild-type and *Zbp1*^fl/fl^ BMDMs (Figure 5A–B). *Zbp1*^fl/fl^*LysM*^cre^ BMDMs were protected from ethanol-induced cell lysis compared to *Zbp1*^fl/fl^ BMDMs, as cell death was reduced from 81% to 48%, while *Zbp1*^−/−^ cells did not exhibit robust cell death by 5 h post-treatment (Figure 5A–B). The activation status of PANoptotic proteins in response to ethanol exhibited a similar pattern as the other triggers. Within 4 h, ethanol-treated wild-type and *Zbp1*^fl/fl^ BMDMs exhibited caspase and executioner activation (Figure 5C). Activation of these PANoptosis molecules was reduced in *Zbp1*^fl/fl^*LysM*^cre^ BMDMs, commensurate with the degree of cell death observed. The low level of activation of these proteins in *Zbp1*^fl/fl^*LysM*^cre^ BMDMs was also consistent with residual expression of ZBP1 (Figure 5C–D), as previously observed with IAV (Figure 3C–D) and KPT-330 plus IFN-γ (Figure 4C–D). These results further demonstrate that *Zbp1*^fl/fl^*LysM*^cre^ BMDMs are resistant to ZBP1-dependent PANoptosis.

**Figure 5.**
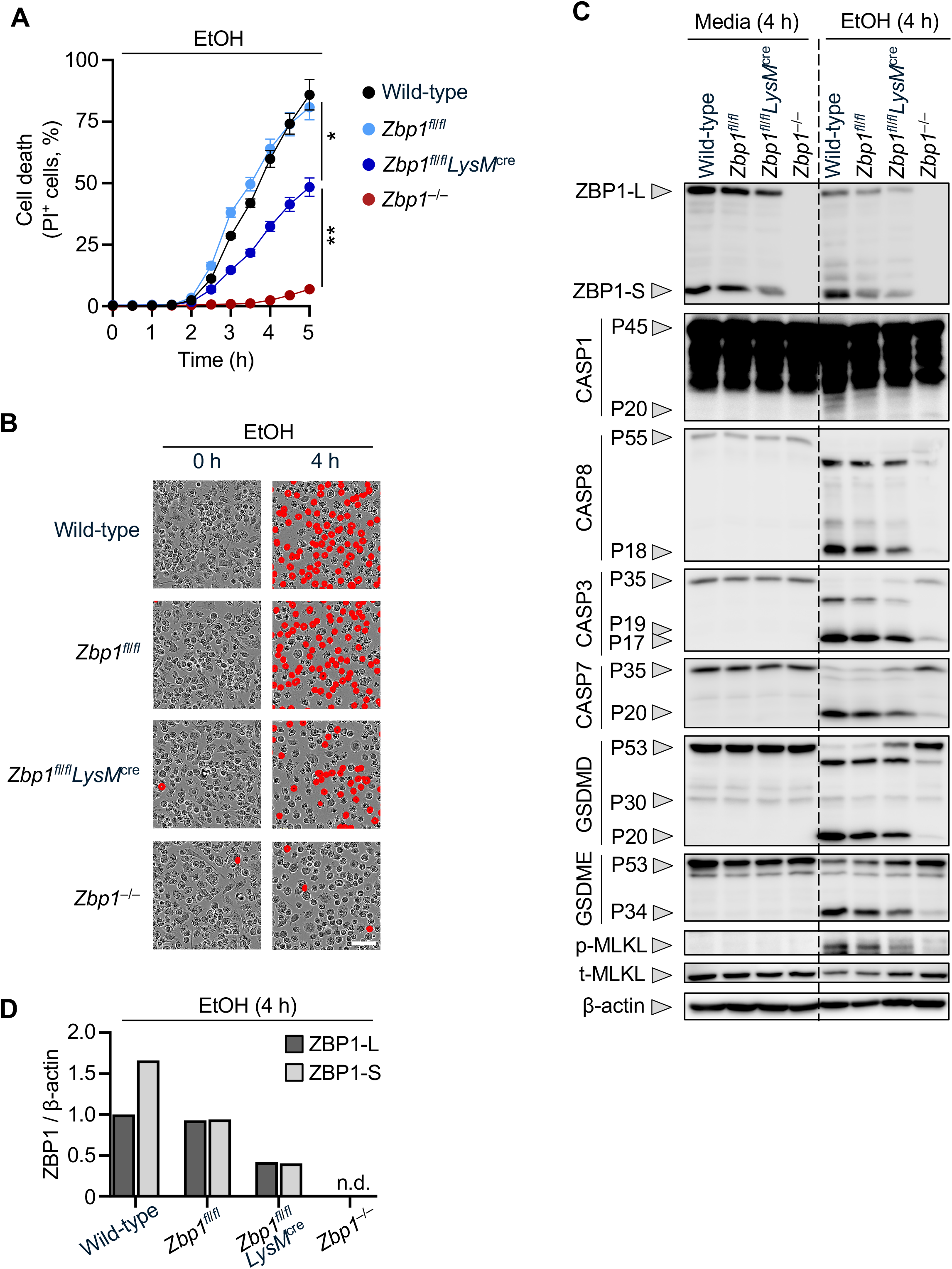
*Zbp1*^fl/fl^*LysM*^cre^ BMDMs undergo reduced PANoptosis in response to EtOH. **A**) Cell death kinetics as assessed by PI uptake in wild-type, *Zbp1*^fl/fl^, *Zbp1*^fl/fl^*LysM*^cre^, and *Zbp1*^−/−^ BMDMs during treatment with ethanol (EtOH, 5% v/v) over 5 h. **B**) Representative live-cell images captured at 0 h and 4 h during treatment with EtOH. Cells that were determined to be PI^+^ are masked in red. **C**) Assessment of cell death proteins following treatment with media or EtOH for 4 h. Immunoblot was used to assess ZBP1 in the long (ZBP1-L) and short (ZBP1-S) isoform, pro- (P45) and active (P20) caspase-1 (CASP1), pro- (P55) and cleaved (P18) caspase-8 (CASP8), pro- (P35) and cleaved (P19/P17) caspase-3 (CASP3), pro- (P35) and cleaved (P20) caspase-7 (CASP7), pro- (P53), active (P30), and inactive (P20) gasdermin D (GSDMD), pro- (P53) and active (P34) gasdermin E (GSDME), phosphorylated MLKL (p-MLKL), and total MLKL (t-MLKL). β-actin was used as a loading control. **D**) ZBP1-L and ZBP1-S proteins were quantified by densitometry relative to β-actin, using the blots presented in panel C. The abundance of ZBP1 was normalized to the full-length (ZBP1-L) isoform in wild-type cells. Data are representative of at least three independent biological replicates (**A**–**D**). Data are presented as mean ± SEM and comprise four technical replicates from one experiment (**A**). Scale bar, 50 μm (**B**). Significance was determined by two-way ANOVA (**A**). * *p* < 0.05, ** *p* < 0.01.

## Discussion

Cell death is critical in the innate immune response, and thus key mediators of inflammatory cell death are attractive therapeutic targets. Understanding the cell-specific effects of innate immune molecules is essential for designing targeted therapeutic approaches that limit off-target effects. Given the critical role of ZBP1 as an innate immune sensor to drive inflammatory cell death, PANoptosis, in a number of diseases, understanding the specific roles of ZBP1 among different cell types will have impacts on therapeutic development. Here, we generate *Zbp1*^fl/fl^ mice to allow for the conditional deletion of *Zbp1*. As a proof-of-principle, we created *Zbp1*^fl/fl^*LysM*^cre^ mice to selectively deplete ZBP1 from myeloid cells. The presence of the floxed *Zbp1* allele or recombination with *LysM*^cre^ did not affect the relative abundance of immune cell populations, including myeloid cells. We further demonstrate that *Zbp1*^fl/fl^*LysM*^cre^ BMDMs were resistant to ZBP1-mediated PANoptosis among infectious and sterile triggers that have been implicated in infection, cancer therapy, and inflammatory disease^8,9,12,16,17,23^. These results indicate that this novel mouse line is useful to investigate ZBP1-dependent PANoptosis in a cell type-specific manner. This tool will be especially useful *in vivo*, allowing investigators to probe the role of ZBP1 in individual cell types in physiologically relevant settings.

Although some ZBP1 protein expression was retained in *Zbp1*^fl/fl^*LysM*^cre^ BMDMs upon stimulation, we still observed significant protection from ZBP1-mediated PANoptosis in these cells. The level of cell death was proportional to the relative reduction of ZBP1 protein expression. *LysM*^cre^ is commonly used for the conditional deletion of genes of interest from macrophages and other myeloid cell types^29^. The recombination efficiency of *LysM*^cre^ can be > 90–100% in alveolar and peritoneal macrophages but < 40% in splenic marginal zone or red pulp macrophages^30^. Thus, the efficiency of *LysM*^cre^ can vary widely, even among macrophage populations. This is consistent with our observation that the efficiency of ZBP1 deletion was incomplete. These results highlight the challenges associated with the development of macrophage-specific conditional knockout mice but also show that even incomplete deletion can be informative to identify roles for ZBP1 in cell death and inflammation.

We anticipate that the *Zbp1*^fl/fl^ mice generated in this study will be useful to study the function of ZBP1 and its roles in inflammation and cell death in a range of cell and tissue types, when combined with the appropriate Cre drivers. ZBP1 was recently shown to sense oxidized mitochondrial DNA in microglia upon exposure to amyloid-β plaques in a mouse model of Alzheimer’s disease^22^. Although brain-resident microglia are functionally similar to macrophages, selective targeting of these cells requires specialized Cre drivers (e.g., *Cx3cr1, P2ry12, Tmem119*). In a mouse model of heart failure with preserved ejection fraction, ZBP1 activation in cardiomyocytes drives PANoptosis downstream of mitochondrial damage and DNA release^31^. In this case, silencing of *Zbp1 in vivo* also suppressed atrial fibrillation following heart failure. Depletion of ZBP1 from cardiomyocytes could be achieved using *Myh6*^cre^ to further examine similar models. Ultimately, the existing evidence suggests that ZBP1 has critical functions in a range of distinct cell types. Therefore, the use of this mouse line in combination with other Cre drivers would be useful to interrogate the distinct functions of ZBP1 in systems such as the cardiovascular system and the central nervous system, which are understudied with respect to the innate immune system.

Overall, we have generated a *Zbp1*-floxed mouse which can be combined with diverse Cre drivers to investigate the role of ZBP1 in a cell type-specific manner. Given the broad roles of ZBP1 across disease states, this new model system will provide a framework for mechanistically understanding ZBP1 functions in discrete cell types and organ systems. This will facilitate the identification of disease mechanisms with critical relevance for therapeutic development.

### Experimental procedures

#### Mice

Wild-type (C57/BL6) and *Zbp1*^−/−32^ mice have been described previously. *Zbp1*-floxed mice (*Zbp1*^fl/fl^) were generated at the Center for Advanced Genome Engineering (CAGE) at St. Jude Children’s Research Hospital using CRISPR/Cas9 technology as described below. All mice were generated on or extensively backcrossed to the C57BL/6 background. *Zbp1*^fl/fl^ mice were crossed to *LysM*^*cre*^ mice (Jackson Laboratory, 004781) to generate *Zbp1*^fl/fl^*LysM*^*cre*^ mice and Cre-negative *Zbp1*^fl/fl^ littermate controls. Mice were bred at the Animal Resources Center at St. Jude Children’s Research Hospital and maintained under specific pathogen-free conditions. Male and female 6-to 12-week-old mice were used for this study. All animal use was conducted under protocols approved by the St. Jude Children’s Research Hospital committee on the Use and Care of Animals. Mice were maintained under specific pathogen-free conditions at the Animal Resources Center at St. Jude Children’s Research Hospital. Mice were fed standard chow and subjected to a 12 h light/dark cycle.

#### Generation of the *Zbp1*-floxed mouse strain

*Zbp1*^fl/fl^ mice were generated using CRISPR/Cas9 technology in collaboration with St. Jude’s Center for Advanced Genome Engineering (CAGE) and the Transgenic/Gene Knockout Shared Resource facilities. Pronuclear-staged C57BL/6J zygotes were injected with Cas9 protein combined with guide RNAs (gRNAs) with loxP single-stranded oligodeoxynucleotides (ssODNs) targeting regions flanking exons 2 and 3 of the *Zbp1* allele. The zygotes were surgically transplanted into the oviducts of pseudo-pregnant CD1 females, and newborn mice carrying the 5’ and 3’ loxP sites flanking exons 2 and 3 were identified by next generation sequencing (NGS) and Sanger sequencing methods. For routine genotyping, both the 5’ junction and 3’ junction were PCR amplified by using the following primers: P1, 5’-CCATGCCTTCGATGTGGGTGCTGAGC-3’; P2, 5’-GCCTAGTCCCCAGAGCAGAGAAGGCA-3’; P3, 5’-GCCAGTGCTCTACCCAGTCAACCACG-3’; P4, 5’-ACATCGTTTTCAGACATGGCTGTGTC-3’ (Figure 1A–C). The details of the generation of the CRISPR reagents were described previously^33^. The uniqueness of gRNAs and the off-target sites with fewer than three mismatches were found using the Cas-OFFinder algorithm^34^.

#### Bone marrow-derived macrophage (BMDM) generation

Primary mouse BMDMs from wild-type and indicated mutant mice were prepared as previously described^35^. Briefly, bone marrow was isolated from the tibia and femurs of 6-to 12-week old mice and grown in 15 cm dishes for 6 days in 25 mL BMDM media (IMDM [12440053, Thermo Fisher Scientific] supplemented with 10% heat-inactivated fetal bovine serum [HI-FBS; S1620, Biowest], 30% L929 conditioned media, 1% non-essential amino acids [11140-050, Thermo Fisher Scientific], and 1% penicillin and streptomycin [15070-063, Thermo Fisher Scientific]). Plates were supplemented with an additional 8 to 10 mL of BMDM media on days 3 and 5, and cells were collected on the sixth day. Upon collection, BMDMs were dispensed into 12-well plates at a density of 1 × 10^6^ cells/well or 24-well plates at a density of 5 × 10^5^ cells/well and incubated overnight to facilitate adherence prior to treatment.

#### Cell infection and treatment

For IAV infection, IAV (A/Puerto Rico/8/34, H1N1 [PR8]) was generated as described previously^8^ and propagated from 11-day-old embryonated chicken eggs by allantoic inoculation. IAV titer was measured by plaque assay in Madin–Darby canine kidney cells (CCL-34, ATCC). BMDMs were infected with IAV at a multiplicity of infection (MOI) of 25 in serum- and glutamine-free DMEM (D5671, Sigma) in a total volume of 225 μL in a 24-well plate or 450 μL in a 12-well plate. After 1 h of infection, HI-FBS was added to a final composition of 10%. For western blotting, sample lysates were collected after 16 h.

For KPT-330 plus IFN-γ treatment, BMDMs were stimulated as described previously^9^. Briefly, KPT-330 (S7252, Selleckchem; 5 μM) and IFN-γ (315-05, Peprotech; 50 ng/mL) were prepared in DMEM containing 10% HI-FBS. BMDMs were rinsed in warm PBS and the solution of KPT-330 plus IFN-γ was applied in a volume of 250 μL for a 24-well plate or 500 μL for a 12-well plate. For western blotting, sample lysates were collected after 24 h.

For ethanol treatment, BMDMs were stimulated as described previously^12^. Briefly, ethanol (04-355-451, Thermo Fisher Scientific; 5% vol/vol) was prepared in DMEM containing 10% HI-FBS. BMDMs were rinsed in warm PBS, and the ethanol solution was applied in a volume of 250 μL for a 24-well plate or 500 μL for a 12-well plate. For western blotting, sample lysates were collected after 4 h.

#### Flow cytometry

Mice were euthanized by CO_2._ Peritoneal lavage samples were obtained from *Zbp1*^fl/fl^*LysM*^cre^ mice and a pool of control mice (wild-type, *Zbp1*^fl/fl^, and *Zbp1*^+/+^*LysM*^cre^ mice) by intraperitoneal injection and subsequent aspiration of 5 mL PBS. A single cell suspension of splenocytes was prepared by mechanical dissociation of the spleen in 5 mL PBS, followed by red blood cell lysis (with the following buffer: 155 mM NH_4_Cl, 10 mM KHCO_3_, 0.1 mM EDTA at pH 7.2) then washed with wash buffer (PBS containing 2% HI-FBS). The cells were stained in wash buffer with antibodies (1:500 dilution) on ice for 30 min, then DAPI (40043, Biotium; 1 μg/mL) was added for an additional 10 min before washing. Cells were Fc-blocked using anti-CD16/32 (Clone 93, Biolegend, 101302) alongside staining with the following antibodies: anti-CD45.2-PE/Cy7 (Clone 104, Biolegend, 109830), anti-CD4-PerCP/Cy5.5 (Clone RM4-5, Biolegend, 100540), anti-CD8-FITC (Clone53-6.7, Biolegend, 100706), anti-CD19-APC/Cy7 (Clone 1D3, Biolegend, 152411), anti-Ly6G-FITC (Clone 1A8, Biolegend, 127605), anti-Ly6C-APC (Clone HK1.4, Biolegend, 128016), anti-F4/80-PE (Clone BM8.1, Tonbo, 50-4801-U100), anti-NK1.1-PerCP/Cy5.5 (Clone PK136, Tonbo, 65-5941-U025), anti-cKit-AF488 (Clone 2B8, Biolegend, 105816), anti-FcεRI-APC (Clone MAR-1, Biolegend, 134316) and anti-SiglecF-PE (Clone E50-2440, BD, 552126). Flow cytometry data were collected using an LSRFortessa system (BD Bioscience) and data were analyzed using FlowJo software (BD Bioscience).

#### Kinetic analysis of cell death

BMDMs were seeded in 24-well tissue culture plates and treated as indicated. Cell death was inferred by uptake of propidium iodide (PI, P3566, Life Technologies; 500 ng/mL) which was monitored using an IncuCyte S3/SX5 (Sartorius) live-cell imaging and analysis system. Plates were scanned to capture phase-contrast images and red fluorescence either every 30 min or 60 min. PI-positive cells were masked (outline, 10; sensitivity threshold, 0.5) and quantified using the IncuCyte live-cell analysis software (Sartorius).

#### Immunoblot analysis

Immunoblotting was performed as previously described^36^. To assess caspase cleavage, caspase lysis buffer (10% NP-40, 25 mM DTT, protease inhibitor [11697498001, Roche] and phosphatase inhibitor [04906837001, Roche]) and 4× SDS sample loading buffer (with 2-mercaptoethanol) was added to cells and combined supernatants. To assess other intracellular and membrane-bound proteins, cells were rinsed once with cold 1× Dulbecco’s PBS (DPBS; 14190-250, Thermo Fisher Scientific) and lysed in RIPA buffer (50 mM Tris, 150 mM NaCl, 1% NP-40 alternative, 0.1% sodium deoxycholate, 0.1% SDS) and 4× SDS sample loading buffer. Samples were boiled at 100°C for 10 min immediately following collection.

Proteins were resolved on 10–12% polyacrylamide gels and transferred to PVDF membranes (IPVH00010, Millipore) using the Trans-Blot Turbo system (Bio-Rad). Membranes were blocked in 5% skim milk and incubated overnight at 4 °C with gentle rocking in primary antibodies, diluted 1:1000 in 5% milk. The following proteins were probed using single antibodies: ZBP1 (AG-20B-0010, AdipoGen), caspase-1 (AG-20B-0042, AdipoGen), GSDMD (ab209845, Abcam), GSDME (ab215191, Abcam), p-MLKL (37333, Cell Signaling Technology [CST]), and t-MLKL (AP14272B, Abgent). The following caspases were probed using a combination of pro-form and cleavage product-specific antibodies: caspase-3 (9662, CST), cleaved caspase-3 (9661, CST), caspase-7 (9492, CST), cleaved caspase-7 (9491, CST), caspase-8 (4927, CST), and cleaved caspase-8 (8592, CST). Membranes were washed thrice in TBS-T for 10 minutes each and incubated with the appropriate horseradish peroxidase (HRP)-conjugated secondary antibodies, diluted 1:5000 in 5% milk (anti-mouse [315-035-047] and anti-rabbit [111-035-047], Jackson ImmunoResearch Laboratories). Membranes were washed thrice more in TBS-T and imaged using Immobilon® Forte Western HRP Substrate (WBLUF0500, Millipore) or SuperSignal™ West Femto Maximum Sensitivity Substrate (34096, Thermo Fisher Scientific) and an Amersham Imager. Following imaging, membranes were stripped with Restore™ Stripping Buffer (Thermo Fisher Scientific) and re-probed for β-actin (sc-47778 HRP, Santa Cruz; 1:5000) or other proteins as needed. For quantification, densitometric measurements were performed using ImageJ software (version 1.53k, NIH).

#### Statistical analysis

GraphPad Prism 10.0 software was used to perform statistical analyses of kinetic cell death data. To compare kinetic data with three or more groups, two-way analysis of variance (ANOVA) was performed, with the Dunnett post-hoc test to correct for multiple comparisons at each time-point. *P* value < 0.05 was considered statistically significant.

## Acknowledgements

We thank all members of the Kanneganti laboratory for their valuable comments and suggestions during the development of this manuscript. We also thank Kristine Zengeler, PhD, and Rebecca Tweedell, PhD, for their assistance with scientific editing and writing support. Work in the Kanneganti lab is supported by NIH grants AI101935, AI124346, AI160179, and CA253095 and the American Lebanese Syrian Associated Charities to T.-D.K. The content is solely the responsibility of the authors and does not necessarily represent the official views of the National Institutes of Health.

## References

1 Zheng, M. K., R; Vogel, P; Kanneganti, TD Caspase-6 is a key regulator of innate immunity, inflammasome activation and host defense. Cell 181, 674–687.e613 (2020).

2 Lee, S. et al. AIM2 forms a complex with pyrin and ZBP1 to drive PANoptosis and host defence. Nature (2021). 10.1038/s41586-021-03875-8

3 Malireddi, R. K. S. et al. RIPK1 Distinctly Regulates Yersinia-Induced Inflammatory Cell Death, PANoptosis. Immunohorizons 4, 789–796 (2020). 10.4049/immunohorizons.2000097

4 Sundaram, B. et al. NLRC5 senses NAD+ depletion, forming a PANoptosome and driving PANoptosis and inflammation. Cell (2024). 10.1016/j.cell.2024.05.034

5 Sundaram, B. et al. NLRP12-PANoptosome activates PANoptosis and pathology in response to heme and PAMPs. Cell 186, 2783–2801.e2720 (2023). 10.1016/j.cell.2023.05.005

6 Sharma, B. R., Choudhury, S. M., Abdelaal, H. M., Wang, Y. & Kanneganti, T.-D. Innate immune sensor NLRP3 drives PANoptosome formation and PANoptosis. J Immunol 214, 1236–1246 (2025). 10.1093/jimmun/vkaf042

7 Christgen, S., Zheng, M., Kesavardhana, S., Karki, R., Malireddi, R.K.S., Banoth, B., Place, D.E., Briard, B., Sharma, B.R., Tuladhar, S., Samir, P., Burton, A., Kanneganti, T.-D. Identification of the PANoptosome: A molecular platform triggering pyroptosis, apoptosis, and necroptosis (PANoptosis). Front Cell Infect Microbiol 10 (2020).

8 Kuriakose, T. et al. ZBP1/DAI is an innate sensor of influenza virus triggering the NLRP3 inflammasome and programmed cell death pathways. Sci Immunol 1 (2016). 10.1126/sciimmunol.aag2045

9 Karki, R. et al. ADAR1 restricts ZBP1-mediated immune response and PANoptosis to promote tumorigenesis. Cell Rep 37, 109858 (2021). 10.1016/j.celrep.2021.109858

10 Karki, R. et al. ZBP1-dependent inflammatory cell death, PANoptosis, and cytokine storm disrupt IFN therapeutic efficacy during coronavirus infection. Sci Immunol 7, eabo6294 (2022). 10.1126/sciimmunol.abo6294

11 Yang, B. et al. SARS-CoV-2 infection induces ZBP1-dependent PANoptosis in bystander cells. Proc Natl Acad Sci U S A 122, e2500208122 (2025). 10.1073/pnas.2500208122

12 Qin, Q. et al. Identification of an IRF–ZBP1–caspase-8–NINJ1 axis in driving PANoptosis and pathology during alcohol-associated liver disease. Proc Natl Acad Sci U S A 122, e2525296122 (2025). 10.1073/pnas.2525296122

13 Wang, S. et al. Fn-OMV potentiates ZBP1-mediated PANoptosis triggered by oncolytic HSV-1 to fuel antitumor immunity. Nat Commun 15, 3669 (2024). 10.1038/s41467-024-48032-7

14 Cui, Y. et al. Ischemia-reperfusion injury induces ZBP1-dependent PANoptosis in endothelial cells. Biochim Biophys Acta Mol Basis Dis 1871, 167782 (2025). 10.1016/j.bbadis.2025.167782

15 Gong, T. et al. Mechanism of lactic acidemia-promoted pulmonary endothelial cells death in sepsis: role for CIRP-ZBP1-PANoptosis pathway. Mil Med Res 11, 71 (2024). 10.1186/s40779-024-00574-z

16 Jiao, H. et al. ADAR1 averts fatal type I interferon induction by ZBP1. Nature 607, 776–783 (2022). 10.1038/s41586-022-04878-9

17 de Reuver, R. et al. ADAR1 prevents autoinflammation by suppressing spontaneous ZBP1 activation. Nature 607, 784–789 (2022). 10.1038/s41586-022-04974-w

18 Wang, F. et al. Sensing of endogenous retroviruses-derived RNA by ZBP1 triggers PANoptosis in DNA damage and contributes to toxic side effects of chemotherapy. Cell Death Dis 15, 779 (2024). 10.1038/s41419-024-07175-7

19 Zhao, Q. et al. Reduction of D2 receptors on microglia leads to ZBP1-mediated PANoptosis of mPFC in Parkinson’s disease depression mice. Int Immunopharmacol 158, 114809 (2025). 10.1016/j.intimp.2025.114809

20 Yang, H. et al. ADAR1 prevents ZBP1-dependent PANoptosis via A-to-I RNA editing in developmental sevoflurane neurotoxicity. Cell Biol Toxicol 40, 57 (2024). 10.1007/s10565-024-09905-1

21 Ren, L. et al. CDK1 serves as a therapeutic target of adrenocortical carcinoma via regulating epithelial–mesenchymal transition, G2/M phase transition, and PANoptosis. J Transl Med 20, 444 (2022). 10.1186/s12967-022-03641-y

22 Song, Z. et al. Innate immune sensing of Z-nucleic acids by ZBP1-RIPK1 axis drives neuroinflammation in Alzheimer’s disease. Immunity 58, 2574–2592 e2579 (2025). 10.1016/j.immuni.2025.07.024

23 Hubbard, N. W. et al. ADAR1 mutation causes ZBP1-dependent immunopathology. Nature 607, 769–775 (2022). 10.1038/s41586-022-04896-7

24 Kesavardhana, S. et al. The Zα2 domain of ZBP1 is a molecular switch regulating influenza-induced PANoptosis and perinatal lethality during development. J Biol Chem 295, 8325–8330 (2020). 10.1074/jbc.ra120.013752

25 Kesavardhana, S. et al. ZBP1/DAI ubiquitination and sensing of influenza vRNPs activate programmed cell death. J Exp Med 214, 2217–2229 (2017). 10.1084/jem.20170550

26 Nagata, M., Carvalho Schafer, Y., Wachsmuth, L. & Pasparakis, M. A shorter splicing isoform antagonizes ZBP1 to modulate cell death and inflammatory responses. EMBO J 43, 5037–5056 (2024). 10.1038/s44318-024-00238-7

27 Cai, Z. Y. et al. A ZBP1 isoform blocks ZBP1-mediated cell death. Cell Rep 43, 114221 (2024). 10.1016/j.celrep.2024.114221

28 Malireddi, R. K. S. et al. ZBP1 Drives IAV-Induced NLRP3 Inflammasome Activation and Lytic Cell Death, PANoptosis, Independent of the Necroptosis Executioner MLKL. Viruses 15 (2023). 10.3390/v15112141

29 Clausen, B. E., Burkhardt, C., Reith, W., Renkawitz, R. & Forster, I. Conditional gene targeting in macrophages and granulocytes using LysMcre mice. Transgenic Res 8, 265–277 (1999). 10.1023/a:1008942828960

30 Abram, C. L., Roberge, G. L., Hu, Y. & Lowell, C. A. Comparative analysis of the efficiency and specificity of myeloid-Cre deleting strains using ROSA-EYFP reporter mice. J Immunol Methods 408, 89–100 (2014). 10.1016/j.jim.2014.05.009

31 Duan, J. et al. Mitochondrial dysfunction drives ZBP1-mediated PANoptosis to increase the susceptibility of heart failure with preserved ejection fraction-associated atrial fibrillation. J Adv Res (2025). 10.1016/j.jare.2025.09.016

32 Ishii, K. J. et al. TANK-binding kinase-1 delineates innate and adaptive immune responses to DNA vaccines. Nature 451, 725–729 (2008). 10.1038/nature06537

33 Pelletier, S., Gingras, S. & Green, D. R. Mouse genome engineering via CRISPR-Cas9 for study of immune function. Immunity 42, 18–27 (2015). 10.1016/j.immuni.2015.01.004

34 Bae, S., Park, J. & Kim, J. S. Cas-OFFinder: a fast and versatile algorithm that searches for potential off-target sites of Cas9 RNA-guided endonucleases. Bioinformatics 30, 1473–1475 (2014). 10.1093/bioinformatics/btu048

35 Tweedell, R. E., Malireddi, R. K. S. & Kanneganti, T. D. A comprehensive guide to studying inflammasome activation and cell death. Nat Protoc 15, 3284–3333 (2020). 10.1038/s41596-020-0374-9

36 Han, J. H., Tweedell, R. E. & Kanneganti, T. D. Evaluation of Caspase Activation to Assess Innate Immune Cell Death. J Vis Exp (2023). 10.3791/64308

